# Evolutionary variation in MADS-box dimerization affects floral development and protein stability

**DOI:** 10.1101/2020.03.09.984260

**Authors:** Maria Jazmin Abraham Juarez, Amanda Schrager-Lavelle, Jarrett Man, Clinton Whipple, Pubudu Handakumbura, Courtney Babbitt, Madelaine Bartlett

**Affiliations:** IPICYT; Colorado Mesa University; University of Massachusetts Amherst; Brigham Young University; Pacific Northwest National Laboratory; UMass Amherst; University of Massachusetts, Amherst

**Author notes:** co-first authors (Maria Jazmin Abraham Juarez, Amanda Schrager-Lavelle).

## Abstract

Shifting interactions between MADS-box transcription factors may have been critical in the emergence of the flower, and in floral diversification. However, how evolutionary variation in MADS-box interactions affects the development and evolution of floral form remains unknown. Interactions between B-class MADS-box proteins are variable across the grass family. Here, we test the functional consequences of this evolutionary variability using maize as an experimental system. We found that differential B-class dimerization was associated with subtle, quantitative differences in stamen shape. In contrast, differential dimerization resulted in large-scale changes to protein complex composition and downstream gene expression. Differential dimerization also affected B-class complex abundance, independent of RNA levels. Thus, differential dimerization may affect protein stability. This reveals an important consequence for evolutionary variability in MADS-box interactions, adding complexity to the evolution of developmental gene networks. Our results show that floral development is robust to molecular change, even coding change in a master regulator of development. This robustness may contribute to the evolvability of floral form.

## Introduction

A paradox lies at the heart of floral evo-devo: flower morphology is diverse, but the master regulators specifying floral organ identity are conserved. These master regulators are the floral MADS-box transcription factors. The ABC model of floral development was derived from mutants in *Antirrhinum majus* and *Arabidopsis thaliana*, and explains how floral organs are specified by groups of transcription factors (A-through E-class proteins). All but one of the original ABCDE genes encode MADS-box transcription factors (Krizek and Fletcher, 2005). These MADS-box proteins likely function as tetramers, with tetramer composition determining

DNA-binding and downstream gene regulation. For example, tetramers of B-, C-, and E-class proteins may specify stamen identity, and tetramers of C and E-class proteins may specify carpel identity (Theissen and Saedler, 2001; Theissen et al., 2016). This model is supported by genetic data, *in vitro* characterization of protein-protein interactions, and *in planta* evidence for MADS-box complex formation and DNA-binding by floral quartets, particularly in *A. thaliana* (Theissen et al., 2016). Outside of the eudicots, B-class and C-class function, in particular, is deeply conserved (Kramer, 2019). For example, B-class genes specify stamen and petal identity in every species with functional data, even when petals are highly modified (Kramer, 2019). How, then, can floral organs vary so extensively in form and function, when they are specified by orthologous genes?

The combinatorial assembly of MADS-box protein complexes has been proposed as an answer to this question. Although floral quartets composed of the same protein classes may specify the same floral organs in different lineages, mixing and matching of particular MADS-box proteins in floral quartets may generate floral diversity by regulating different suites of downstream genes. These differences in downstream targets may contribute to evolutionary variation in organ shape and form (Veron et al., 2007; Mondragon-Palomino and Theissen, 2008; Hsu et al., 2015). Evolutionary changes to MADS-box complexes, specifically the emergence of BCE MADS-box complexes, may also have been important in the evolution of the flower itself (Ruelens et al., 2017; Wang et al., 2010). Thus, combinatorial assembly of MADS-box complexes may have been important in floral evolution and diversification.

Although differential MADS-box complex assembly presents an appealing model for explaining floral diversification, the consequences of changing MADS-box protein-protein interactions have not been extensively tested. The model predicts (1) deep conservation of MADS-box complexes like the BCE complex, and (2) that evolutionary changes to floral MADS-box protein-protein interactions should result in changes to gene regulation, and to floral form. Here, we test these predictions using the B-class MADS box protein STERILE TASSEL SILKY EAR1 (STS1, a homolog of PISTILLATA) (Bartlett et al., 2015). In maize, STS1 forms obligate heterodimers with SILKY1 (SI1, a homolog of APETALA3) (Ambrose et al., 2000). We engineered an ancestral variant of STS1 that forms both homodimers and heterodimers with SI1 (Bartlett et al., 2016). We show that this facultative STS1 homodimerization has subtle effects on stamen development, but large effects on downstream gene expression and protein complex assembly. We also show that BCE complexes do form in maize. Lastly, we found that B-class dimerization affected MADS-box complex stability. Our results show that single changes to MADS-box protein-protein interactions can alter floral development, but likely act in concert with other changes in the evolution of flower morphology.

## Results

### STS1-HET and STS1-HOM show subtle differences in localization and function

To explore the effects of B-class hetero-vs. homodimerization, we developed transgenic maize plants that express a version of STS1 that can bind DNA as homodimers. We found that changing the glycine (G) residue at position 81 to aspartic acid (D) reverts STS1 to its most likely ancestral dimerization state - able to form both homodimers and heterodimers with SI1 (Bartlett et al., 2016). Other amino acid residues differ between STS1 and its most likely ancestor. However, we chose to introduce a single change so that we could explicitly test the effects of hetero-vs homodimerization, to the exclusion of other differences between the extant and ancestral proteins. We introduced the critical G81D change into an STS1-yellow fluorescent protein fusion construct (STS1-YFP) that rescues *sts1* (Bartlett et al., 2015), and used it to transform maize. We will refer to the obligate heterodimer construct (*pSTSl::STSl-YFP*) as *STS1-HET*, and to the STS1 homodimer construct (*pSTS1::STS1(G81D)-YFP*) as *STS1-HOM*.

To understand how variable B-class dimerization affects floral development, we used identical crossing schemes to generate lines carrying either *STS1-HET* or *STS1-HOM*, in backgrounds that were segregating either *si1* or *sts1* (Bartlett et al., 2015; Ambrose et al., 2000). In both *si1* and *sts1*, stamen and lodicule (petal homolog) organ identity is lost (Bartlett et al., 2015; Ambrose et al., 2000). We found that both transgenes complemented *sts1*; neither organ identity nor organ number varied between *STS1-HET* and *STS1-HOM* (Fig. 1, Table S1). In contrast, adult flowers in *si1* mutants carrying *STS1-HOM* were indistinguishable from non-transgenic *si1* mutants (Fig. 1). This indicates that *STS1-HOM* can rescue *sts1* mutants and is functional, but can not compensate for the loss of *SI1* function.

To explore differences between STS1-HET and STS1-HOM in more detail, we analyzed the development of complemented *sts1* mutants. We examined early protein localization of both STS1-HET and STS1-HOM using confocal microscopy. We found that STS1-HET was restricted to lodicule and stamen primordia, as expected (Fig. 1 G-H) (Bartlett et al., 2015). In contrast, protein localization was relaxed in STS1-HOM lines, appearing in gynoecia in addition to lodicule and stamen primordia (Fig. 1J). However, this gynoecial localization was not evident in our immunolocalizations using an anti-STS1 antibody (Fig. S1). In contrast to the proteins, *STS1-HET* and *STS1-HOM* RNAs showed similar localization patterns (Fig. S1). This suggests that the subtle localization differences we detected were regulated at the protein level.

In addition to protein localization differences, morphology differed quantitatively between STS1-HOM and STS1-HET flowers. This included differences in anther aspect ratio, such that anthers of STS1-HOM flowers were wider and shorter than those of STS1-HET flowers while they were still developing (t-test p-value = 6.8e-10, Fig. 1K-L, Data Set S1). In contrast, at anthesis, mature anthers of STS1-HOM flowers were longer and thinner than those of STS1-HET flowers (t-test, p-value = 0.003, Fig. 1M-N, Data Set S2). These differences were despite similarities in anther surface area, which we used as a proxy for size (developing anthers: t-test p-value = 0.259, Fig 1K; anthesis: t-test p-value = 0.367, Fig. 1M). Our results suggest that B-class dimerization may affect anther growth dynamics. Similarly, B-class gene expression affects developmental dynamics in *A. thaliana*, where B-class genes are important for both early and late floral organ development (Wuest et al., 2012). Thus, differential B-class dimerization may quantitatively affect anther developmental dynamics.

**Figure 1.**
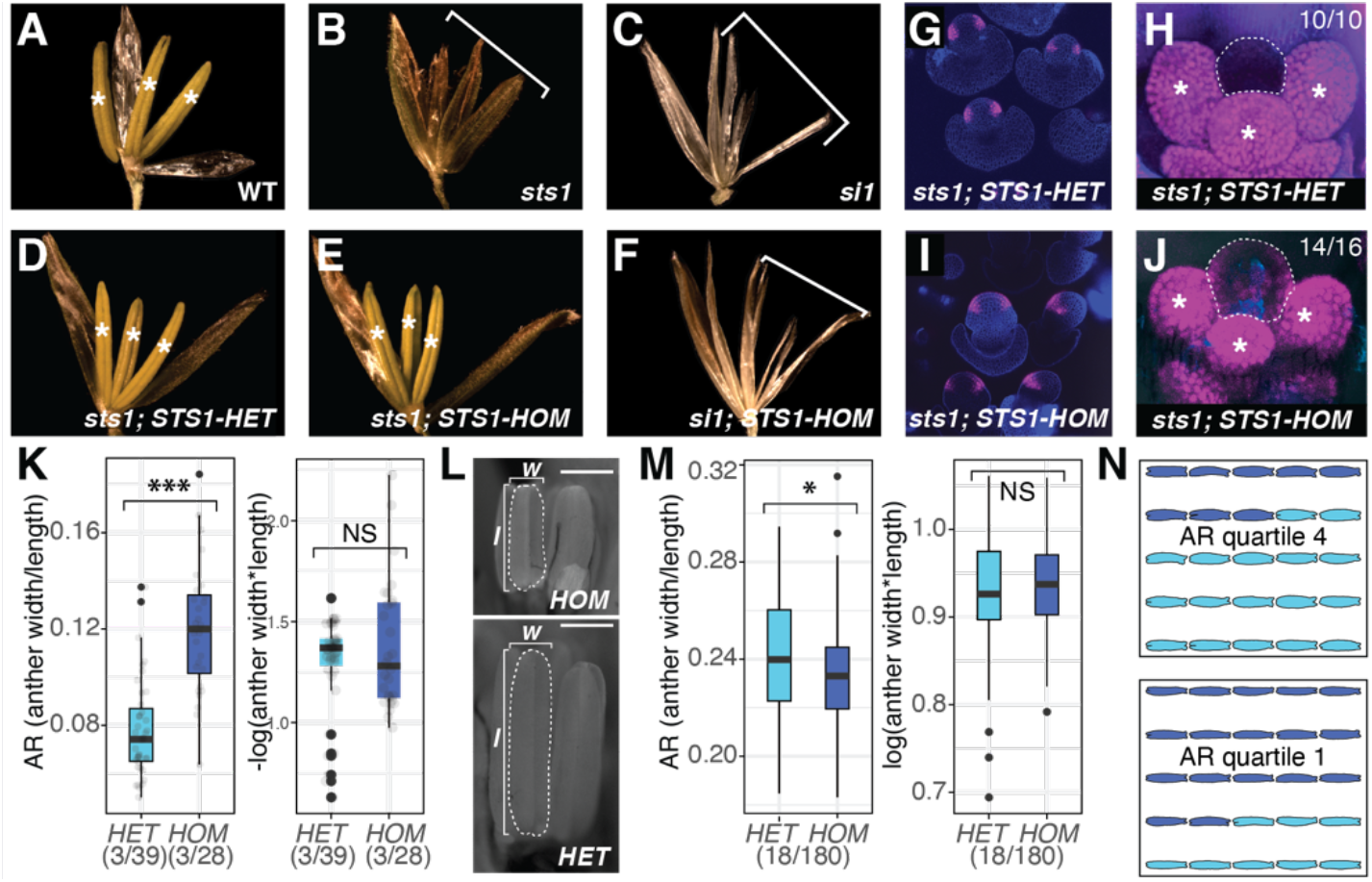
B-class dimerization has subtle effects on floral development in maize. **(A)** Stamen identity (marked with *) is lost in both **(B)** *sts1* **(C)** and *si1* mutant flowers. At anthesis, *sts1* mutant flowers complemented with either the **(D)** *STS1-HET*or **(E)** *STS1-HOM*transgene resemble wild type flowers and each other. (F) The *STS1-HOM* transgene did not complement the si1 mutant phenotype. **(G-J)** Confocal microscopy showing localization of **(G-H)** STS1-HET and (l-J) STS1-HOM localization in developing flowers. Dotted lines in H and J indicate developing gynoecia, numbers in upper right corner indicate frequencies at which we observed the shown localization patterns. **(K-N)** Anther shape metrics **(K-L)** during development and **(M-N)** at anthesis. (K) During development, anther aspect ratio (AR) (anther width/anther length) was higher in *STS1-HOM* anthers than in *STS1-HET*anthers (p-value = p-value = 6.8e-10, left), while anther area (anther width*anther length) was not significantly different (p-value = 0.2592, right). (L) Confocal images of developing anthers measured in **(K)**. **(M)** At anthesis, anther aspect ratio is lower in STS1-HOM anthers than in STS1-HET anthers (p-value = 0.003, left), while anther area is not significantly different (p-value = 0.367, right), p-values calculated using Student’s t-test. **(N)** 25 randomly selected anthers from the first (bottom) and fourth (top) quartiles of anthers measured in M, colored according to STS1 transgene genotype.

### Differential dimerization of maize B-class proteins affects downstream gene regulation

The morphological differences between *STS1-HET* and *STS1-HOM* flowers were subtle, suggesting small phenotypic consequences of differential B-class dimerization. We were curious if molecular function was similarly conserved between *STS1-HET* and *STS1-HOM*. To understand the effect of STS1 dimerization on gene expression, we performed RNA-seq analysis in *sts1* or *si1* mutants complemented with either *STS1-HET* or *STS1-HOM*. We harvested inflorescence tissue shortly after stamen primordium emergence, to capture gene expression just after *STS1* expression initiates (Bartlett et al., 2015). Because genetic diversity is high in maize, we compared expression profiles within genetic backgrounds to control for differences that could arise because of incomplete introgression of the *STS1* transgenes (Buckler et al., 2006). We measured differential expression by comparing expression in each line (*STS1-HOM* and *STS1-HET*) against expression in their mutant siblings.

These analyses revealed more differentially expressed genes in *STS1-HOM* than in *STS1-HET*, as compared to mutant siblings (Fig. 2). At a 5% false discovery rate (FDR), there were 501 differentially expressed genes in inflorescences expressing *STS1-HET*, as compared to *sts1* mutant siblings (Fig. 2A, Data Set S3). In inflorescences expressing *STS1-HOM*, we found 1,257 differentially expressed genes, as compared to *sts1* mutant siblings (Fig. 2B, Data Set S4). There were also 109 shared genes differentially regulated by both *STS1-HOM* and *STS1-HET.* STS1-HOM can both homodimerize and heterodimerize with SI1 (Bartlett et al., 2016). Therefore, to see how gene expression was affected specifically by the STS1 homodimer, we compared gene expression between inflorescences expressing either *STS1-HET* or *STS1-HOM*, in a *si1* mutant background. We found that only 5 genes were differentially expressed in *STS1-HET* inflorescences, as compared to *si1* mutant siblings (Fig. 2C, Data Set S5). In contrast, in inflorescences expressing *STS1-HOM*, 91 genes were differentially expressed, as compared to *si1* mutant siblings (Fig. 2D, Data Set S6). These contrasts indicate that B-class dimerization affects patterns of gene expression in developing inflorescences, either directly or indirectly.

To determine whether the genes regulated by STS1-HET and STS1-HOM were qualitatively similar, we performed GO-term analyses with our differentially expressed gene sets, in an *sts1* mutant background (Data Sets S7 and S8). To compare these lists of enriched GO-terms, we used GO-correlation plots (Fig. 2E-F) (Bergey et al., 2018). Since the *STS1-HET* and *STS1-HOM* constructs were so similar, and since our morphological data suggested subtle functional differences between *STS1-HET* and *STS1-HOM* (Fig. 1), we reasoned that *STS1-HET* and *STS1-HOM* were regulating similar processes. Therefore, we made the threshold for calling a GO-term unique to either dataset very stringent; only GO-terms with an enrichment p-value of less than 0.01 in one dataset, and an enrichment p-value of more than 0.25 in the other dataset were called ‘unique’ (Fig. 2E, sectors i and v). Using these comparisons, we found many GO-terms related to development shared between *STS1-HET* and *STS1-HOM* (Figure 2E). Indeed, 29 of the 65 enriched GO-terms shared between *STS1-HET* and *STS1-HOM* were related to development (p-value in both datasets <0.01, Fig. 2E, sector iii). GO-terms related to signaling and metabolism were also enriched in both datasets, although more of these terms were specifically enriched in *STS1-HOM*. However, the most highly enriched GO-terms in *STS1-HOM* were not significantly enriched in *STS1-HET* (p-value > 0.25). These GO-terms, specific to *STS1-HOM*, were almost all related to chromatin assembly and protein modification (Fig. 2E, sector i). Thus, core floral developmental programs were activated in inflorescences expressing *STS1-HOM*, but B-class dimerization also affected the expression of unique sets of genes, particularly genes involved in chromatin assembly and remodeling.

**Figure 2.**
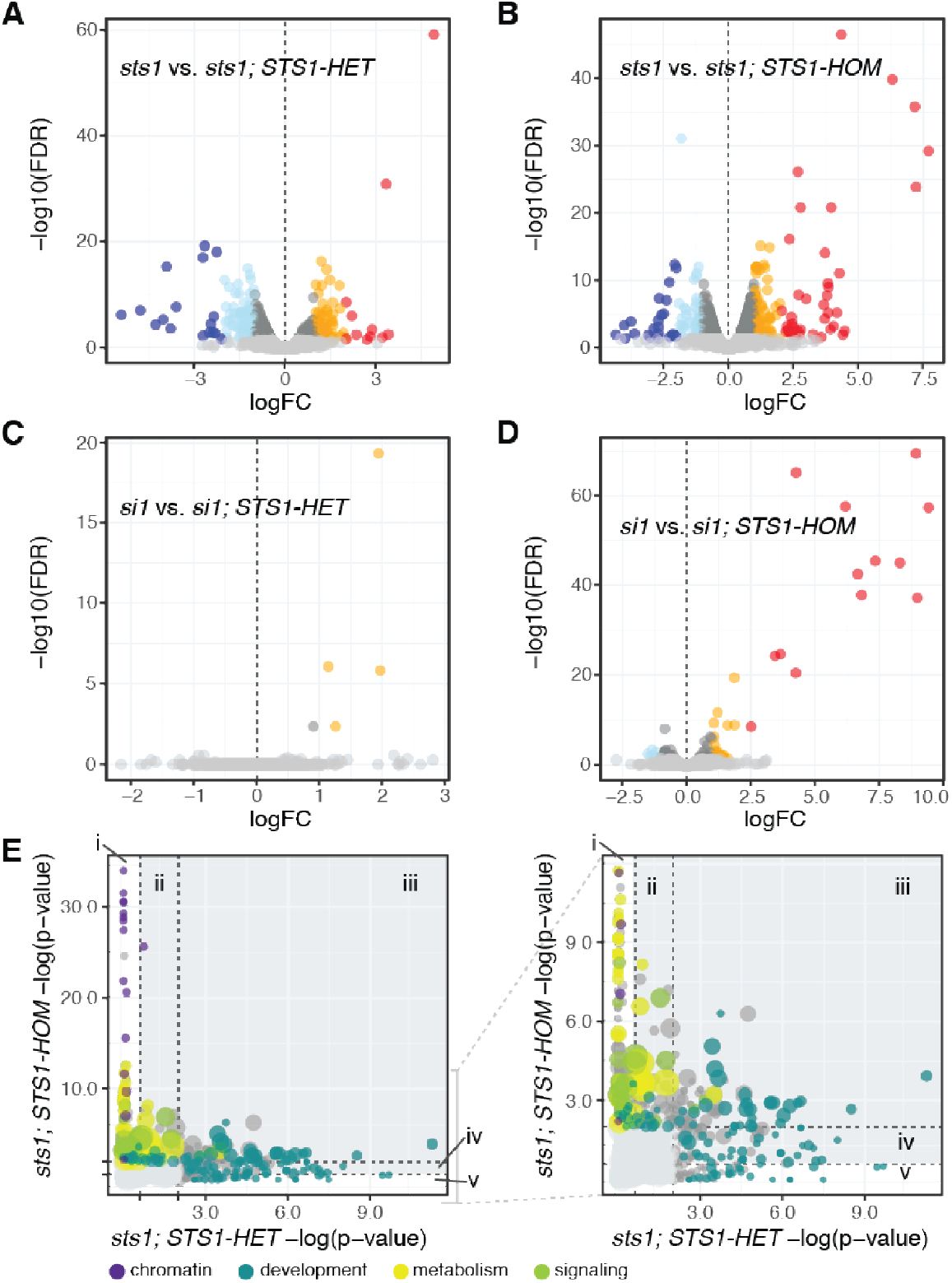
B-class dimerization affects transcriptional profiles in developing tassels. **(A-B)** Differential gene expression in *sts1* mutants complemented with either the **(A)** *STS1-HET*or **(B)** *STS1-HOM* transgene, as compared to *sts1* mutant siblings. **(C-D)** Differential gene expression in *si1* mutants complemented with either the **(C)** *STS1-HET* **(D)** or *STS1-HOM* transgene, as compared to *si1* mutant siblings. (E) GO-term correlation plots comparing probabilities of GO-term enrichments in *STS1-HET*and *STS1-HOM*. Left panel shows all GO-terms, right panel excludes highly enriched GO-terms in *STS1-HOM*. Dot sizes are proportional to the number of genes in each enriched GO-term category, and colored according to which larger category they are associated with. P-value cutoffs for sectors: (i) p-value x > 0.25 & p-value y < 0.01; (ii) 0.01 < p-value x < 0.25 & p.value y < 0.01; (Hi) p-value x < 0.01 & p-value y < 0.01; (iv) p.value x < 0.01 & 0.01 < p-value y < 0.25; (v) p-value x < 0.01 & p-value x > 0.25.

### A complex of B-, C- and E-class proteins is conserved in maize

Differential dimerization could affect the composition of protein complexes, which is crucial for MADS-box function (Theissen and Saedler, 2001; Theissen et al., 2016). To determine which proteins interacted with STS1-HET vs. STS1-HOM, we performed immunoprecipitations (IPs) of STS1-HET and STS1-HOM in an *sts1* mutant background, using a specific antibody against GFP (ChromoTek). We analyzed precipitated complexes using quantitative mass spectrometry (MS). After confirming the presence of STS1 in the IP complex through immunoblotting, we performed trypsin digestion and LC-MS/MS followed by label-free protein quantification (Sinitcyn et al., 2018). The *sts1* mutant was used as a negative control to detect non-specific proteins. The iBAQ (intensity Based Absolute Quantification) method was used to compare the abundances of identified proteins (He et al., 2019; Krey et al., 2014). Protein identification was based on at least 5 exclusive peptides and two replicates were performed for each sample; protein abundances were similar between replicates (Data Sets S9 and S10). We found a higher number of proteins in complex with STS1-HOM; 451 proteins were either specific to STS1-HOM, or were at least 2 fold higher than in mutant siblings (Data Set S9). In contrast, we found 414 proteins in complex with STS1-HET, with the same parameters (Data Set S10).

The first set of proteins in our immunoprecipitations that we explored were the MADS-box proteins. Ancestral protein resurrection and *in vitro* surveys of protein-protein interactions predict that complexes of B-, C-, and E-class MADS-box proteins are conserved across flowering plants (Zhang et al., 2018; Theissen et al., 2016; Veron et al., 2007). However, this prediction remains largely untested *in planta*, especially in monocots. Therefore, we specifically searched for other MADS-box proteins in our IP-MS data. In the STS1-HET IP-MS results, we found both STS1 and SI1 peptides, as well as peptides for three E-class MADS-box proteins (ZmMADS27, ZmMADS7, ZmMADS6), one C-class protein (an AGAMOUS (AG) homolog, ZAG1) and one D-class protein (an AGL5 homolog, ZAG4) (Table S2) (Dreni et al., 2007; Liljegren et al., 2000). In STS1-HOM, we found the same three E-class proteins and the C-class protein ZAG1, but did not find AGL5. AGL5 was identified as one of the most enriched proteins in the STS1-HET IP, and was the only MADS-box protein specific to STS1-HET. Homologs of AGL5 are involved in gynoecium and ovule development in *A. thaliana* and rice, and AGL5 is a direct target of AGAMOUS (Colombo et al., 2010; Dreni et al., 2007; Liljegren et al., 2000). The AGL5 *A. thaliana* homolog, SHATTERPROOF2, is found in complex with AG, but has not been found in complex with PISTILLATA (Smaczniak et al., 2012). This suggests differences in B-class complexes between *A. thaliana* and maize. However, our results indicate broad conservation of B-, C-, and E-class protein complexes, as predicted by models of floral development and evolution (Theissen et al., 2016; Mondragon-Palomino and Theissen, 2008; Ruelens et al., 2017).

### B-class dimerization affected protein abundance

Except for SI1, the MADS-box proteins we identified in both IP datasets were at much higher levels in STS1-HOM than in STS1-HET. This included STS1-HOM itself: absolute iBAQ values for STS1-HOM were 4.0 times higher than for STS1-HET, 3.58 times higher normalized to SI1 (Fig. 3). The higher abundance of STS1-HOM in our immunoprecipitations could have been due to complex stoichiometry, where STS1-HOM homodimerization increased because of double the number of STS1-HOM proteins in MADS-box complexes. STS1-HOM protein levels could also have been generally higher in developing flowers. We suspected higher values overall because our immunolocalizations suggested a higher abundance of STS1-HOM vs. STS1-HET, despite the same experimental conditions (Fig. S1). Because immunolocalizations are not quantitative, we carried out semi-quantitative immunoblots using a polyclonal antibody against STS1 in STS1-HOM vs. STS1-HET inflorescence tissue. In these blots, STS1-HOM was seven times more abundant than STS1-HET (Fig. 3B, Table S3). Together with our IP-MS results, these data indicate that STS1-HOM accumulated to a higher abundance than STS1-HET in inflorescence tissue.

This unexpected protein level difference led us to explore what mechanism could be responsible. STS1-HOM protein could be more abundant than STS1-HET protein because *STS1-HOM* is expressed at higher levels than *STS1-HET*. To test for this possibility, we carried out RT-qPCR using specific primers for *STS1* and *SI1* in *sts1* mutants complemented with either *STS1-HET* or *STS1-HOM*. We found that *SI1* was expressed to the same level in both lines; however, the expression of *STS1* in *STS1-HET* was 3.7-fold higher than in *STS1-HOM* (relative to actin, Fig. 3C). Similarly, in our RNA-Seq results, normalized *STS1-HET* expression was consistently double normalized *STS1-HOM* expression; approximately 15,000 counts vs. 8,600 counts respectively. Thus, despite relatively low *STS1-HOM* expression, STS1-HOM protein accumulates to higher levels than STS1-HET in floral tissue. This suggests that STS1 homodimerization led to increased protein accumulation independent of RNA levels.

**Figure 3.**
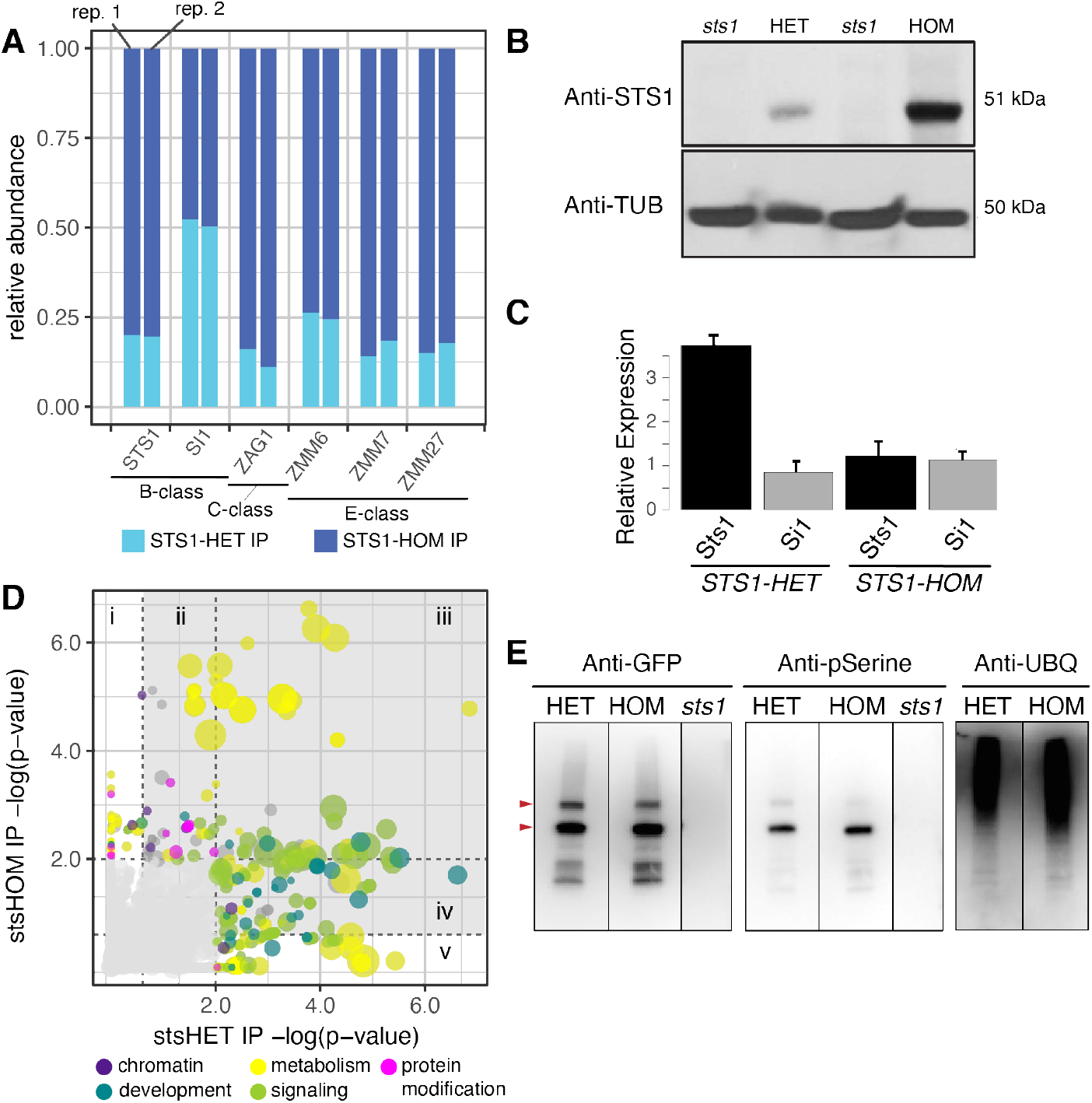
B-class dimerization affects protein abundance and complex assembly in developing tassels. **(A)** Relative abundances of MADS-box proteins in the IP-MS datasets. STS1, as well as C-class and E-class proteins are higher in STS1-HOM than in STS1-HET immunoprecipitations (IPs). **(B)** Semi-quantitative immunoblots with anti-STS1 (top) and anti-TUBULIN (bottom) also show that STS1-HET is less abundant than STS1-HOM. **(C)** RT-qPCR shows that *STS1-HET*RNA is more abundant than *STS1-HOM* RNA, relative to *actin. SI1* RNA occurs at similar levels, relative to *actin,* in both *STS1-HET*and *STS1-HOM*. **(D)** GO-term correlation plot comparing probabilities of GO-term enrichments in the STS1-HET vs. STS1-HOM IP-MS datasets. Dot sizes are proportional to the number of genes in each enriched GO-term category, and colored according to which larger category they are associated with, p-value cutoffs for sectors: (i) p-value x > 0.25 & p-value y < 0.01; (ii) 0.01 < p-value x < 0.25 & p.value y < 0.01; (Hi) p-value x < 0.01 & p-value y < 0.01; (iv) p.value x < 0.01 & 0.01 < p-value y < 0.25; (v) p-value x < 0.01 & p-value x > 0.25. (E) Immunoblots with anti-pSer and anti-Ubiquitin show that STS1-HET and STS1-HOM are phosphorylated, and in complex with ubiquitylated proteins.

The other MADS-box proteins that we detected in our immunoprecipitations were also more abundant in STS1-HOM than in STS1-HET. ZAG1, ZmMADS6 and ZmMADS7 increased in abundance 2-to 9-fold in STS1-HOM, as compared to STS1-HET (Fig. 3A). As with STS1-HOM, higher amounts of ZAG1, ZmMADS6, ZmMADS7, and ZmMADS27 could have been because of differences in protein complex assembly or persistence in the presence of STS1-HET vs. STS1-HOM. Alternatively, the genes encoding these proteins could have been upregulated by STS1-HOM, resulting in higher protein abundance. To distinguish between these possibilities, we looked for these genes in our RNA-Seq data, and found that they were not differentially expressed between *STS1-HOM* and their mutant siblings. From all the identified MADS-box proteins, SI1 was the only one found at similar levels in both STS1-HET and STS1-HOM immunoprecipitations. This suggests that SI1 is able to compete with STS1-HOM to form heterodimers and that SI1 may be a limiting factor in MADS-box protein complex assembly. Taken together, our results show that STS1 homodimerization is either directly affecting the assembly of MADS-box complexes, or indirectly affecting complex persistence because of its higher abundance.

### STS1-HET and STS1-HOM form complexes with chromatin remodelers, kinases and the ubiquitylation machinery

To explore our IP-MS datasets further, we performed GO-term enrichment analyses with identified proteins in the STS1-HET vs. STS1-HOM immunoprecipitations. In these analyses, we only included proteins with at least a two-fold change as compared to *sts1* siblings. When we compared the resulting lists of enriched GO-terms, our results were similar to the RNA-Seq comparisons: development, signaling, and metabolism-related GO-terms were enriched in both datasets to varying degrees. However, chromatin-related GO-terms were no longer exclusively enriched in the STS1-HOM dataset (Fig. 2E, 3A, Data Sets S11 and S12).

We found GO-categories related to chromatin modification enriched in both the STS1-HOM and the STS1-HET immunoprecipitation (Fig. 3D). One specific protein related to chromatin modification was the ISWI chromatin-remodeling complex ATPase CHR11 (Gene ID Zm00001d040831), which we found in complex with both STS1-HET and STS1-HOM. One protein we found only in the STS1-HOM immunoprecipitation was a FRIGIDA-like protein (Zm00001d039594). In *A. thaliana*, FRIGIDA functions as a scaffold for recruiting chromatin modification factors (Hu et al., 2014). MADS-box proteins have long been known to associate with the chromatin remodeling machinery in *A. thaliana*, including with a homolog of CHR11 (Smaczniak et al., 2012; Vachon et al., 2018). Our results show that this may also be the case in maize, and that B-class dimerization may impact associated transcriptional dynamics.

We also found a class of proteins related to signaling in our immunoprecipitation datasets, specifically kinases. We found that ten kinases were immunoprecipitated with STS1-HET, and eleven with STS1-HOM; only two of these were in both samples. Interestingly, the kinases found with STS1-HET were related to basal metabolism, for example a phosphoglycerate kinase (Zm00001d015376) and a pyruvate kinase (Zm00001d023379), involved in synthesis and degradation of carbohydrates (Rosa-Téllez et al., 2018; Lu and Hunter, 2018). In contrast, the kinases that immunoprecipitated with STS1-HOM were related to signaling and membrane receptors; for example BR-signaling kinase 2 (Zm00001d030021) and BR-LRR receptor kinase (Zm00001d052323) (Tang et al., 2008; Xu et al., 2014). However, further analyses are necessary to confirm interaction of STS1 with these kinases.

One new set of GO-terms that emerged in our analysis was related to protein modification and ubiquitylation. These GO-terms were more enriched in STS1-HOM, but were still present in STS1-HET (Fig. 3D). When we explored which proteins might be represented by these enriched GO-terms, we found five proteins of the Cul4-RING E3 ubiquitin ligase complex (CAND1, CUL1, CUL4, XPO1A and CDC) in the STS1-HOM IP, two of which (CAND1 and CUL1) were also in with STS1-HET IP. Although they have not been reported as MADS-box interactors, these proteins are in the same complex in other species, giving us additional confidence in our IP results (Feng et al., 2004; Wertz et al., 2004; Adhvaryu et al., 2015). Thus, our IP-MS results indicate that MADS-box complexes in maize may interact with the ubiquitylation machinery.

### STS1 is post-translationally modified

The presence of kinases and ubiquitylation machinery in our IP results suggested that the higher abundance of STS1-HOM may have been regulated at the protein level by posttranslational modifications. Therefore, we explored the phosphorylation and ubiquitylation of STS1-HET vs. STS1-HOM. To do this, we performed immunoblots using commercial Anti-pSer and Anti-Ubiquitin monoclonal antibodies (Santa Cruz Biotechnology). We found that both STS1-HET and STS1-HOM proteins were phosphorylated (Fig. 3E). When we analyzed IP complexes with an Anti-Ubiquitin antibody, we identified ubiquitylated proteins in both the STS1-HET and the STS1-HOM complex. We detected no differences in phosphorylation or ubiquitylation between STS1-HET and STS1-HOM (Fig. 3E). This indicates that STS1-HET and STS1-HOM are both phosphorylated and in complex with ubiquitylated proteins, and also suggests that the higher abundance of STS1-HOM is not because it is differentially phosphorylated or ubiquitylated. Instead, the higher abundance of STS1-HOM may have allowed us to detect interactions between B-class proteins and the ubiquitylation and phosphorylation machinery that might otherwise have gone unnoticed.

## Discussion

Here, we show that floral development in maize is robust to an evolutionary shift in B-class MADS-box dimerization. This robustness is predicted by network analyses of MADS-box protein-protein interactions, and may be important in the evolution of floral form (Zhang et al., 2018). Very few MADS-box protein-protein interactions have been examined in a comparative framework. However, given evolutionary variation in interactions between B-class proteins (the least promiscuous of the floral MADS-box proteins); the propensity for rapid evolution of MADS-box protein-protein interactions; and lineage-specific MADS-box gene duplications, MADS-box protein-protein interactions profiles are likely diverse within populations and species (Silva et al., 2015; Melzer et al., 2014; Bartlett et al., 2016; Soyk et al., 2017; Alhindi et al., 2017). This cryptic molecular diversity may contribute to the overall evolvability of floral morphology; when genetic variation can accumulate in a system, without consequent phenotypic change, the system itself can be more evolvable (Wagner, 2012). Indeed, morphological floral traits like the lengths of stamens, stigmas, and corolla tubes are evolvable, and can change under selection after only short periods of time (Opedal, 2019; Opedal et al., 2017; Conner et al., 2011). Selection on floral form may also not be limited by genetic variation, indicating underlying variation in the genetic networks that regulate flower development (Ashman and Majetic, 2006). Variation in MADS-box protein-protein interaction networks may represent some of that genetic variation - acting as grist for the mill of natural selection.

MADS-box genes can regulate both complex suites of traits, like organ identity, and individual organ traits, like organ size (Wuest et al., 2012). We found that variation in B-class MADS-box dimerization could affect one aspect of anther shape, namely aspect ratio. Subtle variation in anther shape can impact pollination biology. For example, in the eudicots *Plantago* and *Thalictrum*, anther properties are important for efficient pollen release from anthers (Timerman and Barrett, 2019; Timerman et al., 2014). In *Primula*, another eudicot, variation in anther height is critical in the evolution of heterostyly. Interestingly, a homolog of *STS1* may underlie this height variation(Li et al., 2016). In maize, and likely also in *Primula*, genetics using null mutant phenotypes would have obscured roles for master regulators like the B-class genes in the development of individual organ traits. In contrast, dissecting the evolutionary consequences of small changes in gene sequence can reveal specific functions for pleiotropic genes. This is critical in this era of genome editing, where the technical ability to edit specific DNA base-pairs exists, but the fundamental knowledge of exactly which base-pairs to edit is lacking.

The evolution of MADS-box complex assembly has long been considered only in terms of combinatorial complex assembly. Our results show that evolutionary variability in MADS-box complexes may also affect protein degradation dynamics. Our data suggests that complexes containing STS1 are ubiquitylated and degraded in the proteasome, and that B-class dimerization may affect the dynamics of this degradation. This variable degradation could affect the spectrum of MADS-box complexes present at any one time and, in turn, downstream gene regulation and floral development (Ruelens et al., 2017). In *A. thaliana*, MADS-box proteins are degraded as a result of interactions with effector proteins from phytoplasma pathogens (MacLean et al., 2014). Some phytoplasma effectors have evolved convergently to resemble the protein-protein interaction domains of MADS-box proteins (Aurin et al., 2019; Rümpler et al., 2015). Interactions between MADS-box proteins and these MADS-resembling effectors are likely critical in shuttling MADS-box proteins to the proteasome under phytoplasma infection (Aurin et al., 2019; MacLean et al., 2014; Kitazawa et al., 2017). Our results suggest that phytoplasma effectors may be hijacking a natural process where MADS-box proteins are ubiquitylated and degraded. The effect of MADS-box complex assembly on protein degradation that we uncovered reveals an additional layer of complexity in the evolution of floral development.

## Methods

### Transgenic lines and plant growth

*STS1* transgenes, in the pTF101 vector backbone, were transformed into the Hi-II genetic background at the Iowa State University plant transformation facility. *STS1-HET* and *STS1-HOM* transformants were crossed to either *sts1* or *si1* mutants in the A619 genetic background. Resulting progeny carrying a transgene were identified by their ability to resist herbicide application, and crossed again to *sts1* or *si1* mutants in the A619 genetic background. Once generated, plants for all molecular analyses were grown in the UMass College of Natural Science greenhouse using a 50:50 soil mix of LC1 (SunGro Horticulture) and Turface (Turface Inc.). Field grown plants were grown at the University of Massachusetts UMass Crop and Animal Research and Education Farm in South Deerfield, MA.

### RNAseq tissue collection and sequencing

Plants were grown in the greenhouse as described above. Shortly after stamen primordium emergence (4-5 weeks after planting), plants were harvested and inflorescence meristems were flash frozen in liquid nitrogen. Samples were harvested at the same time of day, beginning at 3pm. Three plants per genotype were pooled to generate one biological replicate, with three biological replicates per genotype (genotyping primers in Table S4). RNA was extracted from each pooled biological replicate using a combination of Trizol (Invitrogen) and Qiagen Plant RNeasy columns including a Qiagen on-column DNase digestion. 1 μg of RNA from each biological replicate was used for RNA library preparation the NEBNext Ultra library kit per manufacturer instructions (3 libraries per genotype). Samples were barcoded using NEBNext Set 1 Multiplex Oligos for Illumina to generate libraries for single end 150 base pair sequencing. DNA sequencing was performed at Genomics Resource Laboratory, University of Massachusetts Amherst using an Illumina NextSeq500.

### Differential expression analysis

Quality and adapter filtering were performed as a part of the Illumina pipeline. Reads were mapped to the version 4 maize genome assembly Zm-B73-REFERENCE-GRAMENE-4.0 using STAR v2.5.3a (Dobin et al., 2013; Jiao et al., 2017). Mapping using STAR included default parameters for alignment and seeding, quality filtering, trimming, and removal of alignments with non-canonical splice junctions to obtain counts per gene (Dobin et al., 2013). Differential expression analysis was performed using the R package RUVseq for normalization, followed by differential expression analysis with edgeR (Robinson et al., 2010; McCarthy et al., 2012; Risso et al., 2014). The RUVseq pipeline included upper quartile normalization using 7,000 empirically determined control genes. These empirical control genes are the 7,000 least differentially expressed genes in the dataset as determined by the analysis pipeline. Where samples did not separate clearly by our experimental variables (e.g. genetic background), those samples were not included in the downstream DE analysis, resulting in the exclusion of one STS1-HET and one STS1-HOM library.

### Confocal microscopy

Plants were grown in the greenhouse as described above. Shortly after stamen primordium emergence (4-5 weeks after planting), plants were harvested and meristems were stained with 5 μM SynaptoRed membrane dye (VWR 80510-682) in DMSO. The confocal image data was gathered using an A1R: Nikon A1 Resonant Scanning Confocal.

### Anther shape measurement and analysis

For measuring anthers and counting floral organs we grew plants in the greenhouse (young anthers) or field (anthers at anthesis) as described above. To measure anthers in young flowers, plants were harvested at the inflorescence meristem stage after glume development (5-6 weeks after planting). Developing flowers were imaged using Leica CTR5500 fluorescent and Zeiss 710 confocal microscopes. Anthers were measured in 3 individuals per genotype, >6 anthers per individual. To measure anthers at anthesis, dehisced anthers were harvested from the central spike on the day that flowers first opened. Anthers with filaments attached were harvested from central spikes and scanned within an hour after dehiscing. Individual anthers were placed on a slide (flat side down) and scanned using an Epson V700 scanner. Scanned images were separated into individual files and images made binary using ImageJ (Schneider et al., 2012). Individual anther image files were read into R, and anther length and width was measured using the R package MOMOCS (Bonhomme et al., 2014). Anther aspect ratio was calculated by dividing anther width by anther length (18 individuals per genotype, 10 anthers per individual). These aspect ratio values were log-transformed (log base 10) for normality (Shapiro-Wilk normality test, p-value = 0.66). Means of log-transformed aspect ratio values were compared using Students-t-test. For the organ counts, mature anthers were harvested from the central spike prior to dehiscence and we counted lodicules and stamens in five spikelets from five individuals (50 florets total). All anther measurements are in supplemental data sets S1 and S2.

### Immunoprecipitation

Five grams (forty plants) of pooled maize tassels (0.5 - 1.0 cm in length) were ground in liquid nitrogen using mortar and pestle. The resulting powder was mixed with 10 mL of an extraction buffer for proteins dynamically transported between nucleus and cytosol, in native conditions (50mM Tris-HCl pH 7.5, 150 mM NaCl, 1% IGEPAL-CA-630 and 1× protease inhibitor mix). Then, extract was filtered through four layers of Miracloth, and centrifuged twice at 10 000rpm for 10 min at 4°C. Protein extract was incubated with 40 uL of GFP-trap MA bead slurry (ChromoTek), shaking for 2 h at 4°C. Beads with the bound target protein were magnetically separated and washed four times with 200 uL of ice-cold wash buffer containing 50 mM Tris pH 7.5, 150 mM NaCl, 0.1% IGEPAL-CA-630 and 1X protease inhibitor mix. Bound proteins were eluted with 40 uL of elution buffer (0.05% Bromophenol blue, 0.1M DTT, 10% Glycerol, 2% SDS, 0.05M Tris-HCl pH 6.8). 5 uL were used for immunoblot to confirm the presence of the bait protein (STS1-YFP) by standard SDS-PAGE and detection by chemiluminescence with a monoclonal Anti-GFP antibody (Mouse IgG1K, clones 7.1 and 13.1, Roche Cat. No. 11814460001) and Anti-Mouse secondary antibody HRP-conjugated (Amersham ECL GE Cat. No. 45001275). Then, 35 uL was used to run a short SDS-PAGE gel, stained with GelCode Blue Safe protein stain (Thermo Fisher scientific). Gel slices were sent for mass spectrometry analysis to UMass Medical School Mass Spectrometry facility. Two replicates were performed for each genotype.

### LC-MS/MS and label-free protein quantification

In-gel trypsin digestion was analyzed in a quadrupole-Orbitrap hybrid mass spectrometer (Thermo Sci Q-Exactive with Waters NanoAcquity UPLC). Forty individuals of each genotype (*sts1* mutants and *sts1* mutants complemented with STS1-HET or STS1-HOM) were used for each IP experiment. When a high number of individuals is used, a lower number of technical replicates in the mass spectrometer is needed to get robust results (Jorrín-Novo et al., 2015). In this experiment the reproducibility can be observed in the high number of proteins identified in both replicates for each sample, as well as in the similar abundance of each one (Supplemental Data Sets S9 and S10). For protein identification, Mascot in Proteome Discoverer 2.1.1.21 with the Uniprot_Maize database was used. For label free quantification, Scaffold version 4.8.4 was used, with 90% minimum peptide threshold, 3 peptides minimum and a peptide FDR 0.05.

### Antibody production and immunolocalization

Anti-STS1 antibody was developed from the fulllength coding sequence of *STS1* cloned into pDEST17 at the Bartlett Lab, using the protocol described in (Chuck et al., 2014) with some modifications. 6HIS-STS1 was expressed and purified from *E. coli* Rosetta strain, using denaturing conditions. 200 ug of purified protein was sent to Cocalico Biologicals, PA., where two guinea pig immunizations were performed. Serum was used for antibody affinity purification to the STS1 recombinant protein using magnetic beads (Invitrogen). Validation of antibody was carried out by immunoblot with total protein extract from *sts1* complemented lines and *sts1* mutant as a negative control. STS1 immunolocalizations were performed as previously described (Tsuda and Chuck, 2019; Chuck et al., 2010).

### STS1 and pSerine immunoblot

SDS-PAGE with 12% acrylamide gel was carried out with 30 ug of protein extract from *sts1*, and STS1-HET and STS1-HOM complemented mutants. Then, semidry transfer, blocking, and incubation with 1:2000 affinity-purified Anti-STS1 guinea pig polyclonal antibody was performed. Protein was detected using chemiluminescence with 1:3000 Anti-Guinea pig secondary HRP-coupled antibody (Thermo Fisher Sci Cat No. A18769). Membrane was stripped and tubulin was detected as a loading control by incubation with 1:25,000 Mouse monoclonal anti-TUB DM1A (Abcam). Detection was done using 1:10,000 Anti-Mouse secondary HRP-conjugated antibody (Amersham ECL GE Cat. No. 45001275). For Anti-pSerine immunoblot, PhosSTOP (Roche) was added to the protein extraction buffer and blocking was carried out with 2% BSA instead milk. Primary Anti-pSer 16B4 mouse monoclonal antibody (Santa Cruz Biotechnology) was used at 1:1000. And secondary Anti-mouse HRP-conjugated (Amersham ECL GE) at 1:3000 was used. For development, Clarity ECL reagents (BioRad) and Azure c-300 Chemiluminescent immunoblot Imaging System (Azure Biosystems) were used.

### In situ hybridization and gel shifts

*sts1* and *sts1* complemented inflorescences were fixed overnight at 4°C in 4% paraformaldehyde in 1X PBS. Fixed samples were dehydrated in ethanol series and transferred into Histoclear, then were embedded in Paraplast, sectioned, and hybridized according to (Bartlett et al., 2015). Gel shifts using B-class proteins were performed as described in (Bartlett et al., 2016).

### Data availability

Raw sequencing data is available at NCBI SRA (link to follow). Supplemental data sets and anther images are available as a dryad data repository (link to follow). Code for analyses and figure generation is available on GitHub (link to follow).

## Author Contributions

MB, AS-L, MJAJ, and CW designed the research. AS-L, MJAJ, MB, CB and JM conducted research and analysed results. MB, AS-L, and PH acquired funding for the research. MB, MJAJ, and AS-L wrote the paper with assistance from co-authors.

## Acknowledgements

We thank Ravi Ranjan at the IALS Genomics Resource Laboratory, and James Chambers at the IALS Light Microscopy Facility and Nikon Center of Excellence. We thank greenhouse and farm staff for essential help: Dan Jones, Chris Joyner, Chris Phillips, and Neal Woodard. We also thank members of the Bartlett lab for help with lab work, and for providing helpful comments on the manuscript: Amanda Dee, Joseph Gallagher, Michelle Heeney, Jeff Heithmar, Harry Klein, Jamie Kostyun, Erin Patterson, and Grace Pisano. Funding was provided by the NSF, the USDA, and the Massachusetts Life Science Center.

## Supplemental Information

### Supplemental Figures

**Figure S1.**
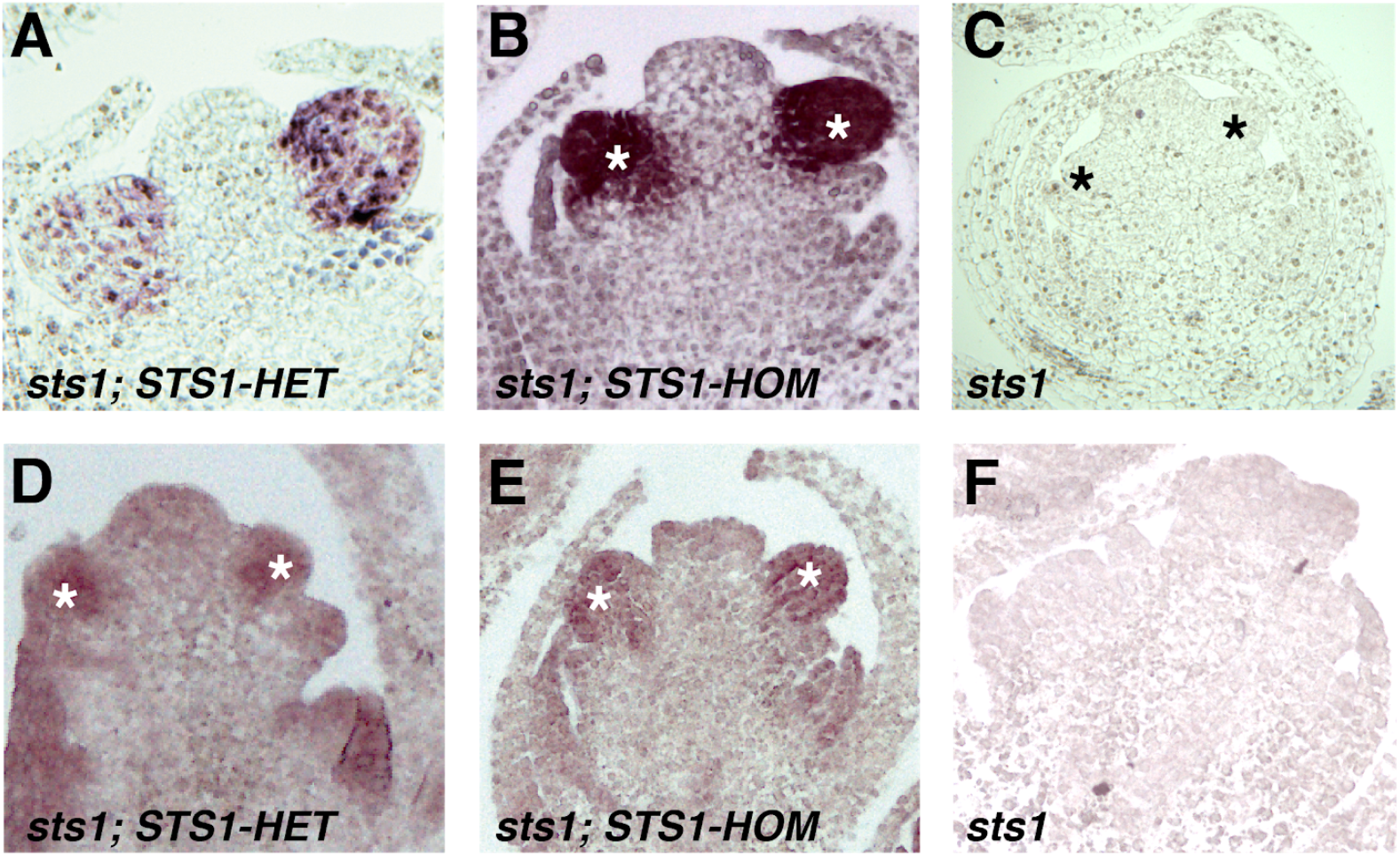
STS1 Immunolocalizations and *in situ* hybridizations. (A-C) STS1 immunolocalizations using a polyclonal anti-STSl antibody in (A) STS1-HET, (B) STS1-HOM and (C) *stsl* mutant flowers. (D-F) *in situ* hybridizations of *STS1* mRNA in (D) *STS1-HET* and (E) *STS1-HOM* (F) *stsl* mutant flowers.

### Supplemental Tables

**Table S1.** Floral organ counts

| **Table S1. Floral organ counts** |  |  |  |  |
| --- | --- | --- | --- | --- |
| **Genotype** | **Number of individuals** | **Florets per plant** | **Stamen number** | **Lodicule number** |
| STS1-HET | 5 | 5 | 2.95 +/- 0.39 | 2.00 +/- 0 |
| STS1-HOM | 5 | 5 | 2.98 +/-0.22 | 2.05 +/- 0.2 |
|  |  |  | NS P=0.57 | NS P=0.08 |

**Table S2.** MADS-box proteins identified in IP-MS experiments

| **Table S2. MADS-box proteins identified in IP-MS experiments** |  |  |  |
| --- | --- | --- | --- |
| **Maize Name** | **Maize ID** | **MADS Clade** | **Arabidopsis homologs** |
| STS1/ZMM16 | Zm00001d042618 | B-class | AT5G20240 |
| SILKY1 | Zm00001d036425 | B-class | AT3G54340 |
| ZMM6 | Zm00001d031620 | E-class | AT3G02310, AT2G03710, AT5G15800, AT1G24260 |
| ZMM7 | Zm00001d021057 | E-class | AT3G02310, AT2G03710, AT5G15800, AT1G24260 |
| ZMM27 | Zm00001d006094 | E-class | AT3G02310, AT2G03710, AT5G15800, AT1G24260 |
| ZAG1 | Zm00001d037737 | C-class | AT3G58780, AT4G18960, AT2G42830 |
| AGL5/ZAG4 | Zm00001d039434 | D-class | AT2G42830 AT4G18960 AT3G58780 |

**Table S3.** Immunoblot quantification of STS1-HET and STS1-HOM

| **Table S3. Immunoblot quantification of STS1-HET and STS1-HOM** |  |  |  |  |
| --- | --- | --- | --- | --- |
| **Genotype** | **STS1** | **TUB** | **Relative intensity** | **Relative intensity to HET** |
| *sts1* | 0 | 30242.43 | 0 | 0 |
| *sts1; STS1-HET* | 17705.066 | 23658.492 | 0.74835987 | 1 |
| *sts1* | 0 | 31542.463 | 0 | 0 |
| *sts1;* | 149610.474 | 25640.48 | 5.834932653 | 7.796960909 |
| *STS1-HOM* |  |  |  |  |

**Table S4.**
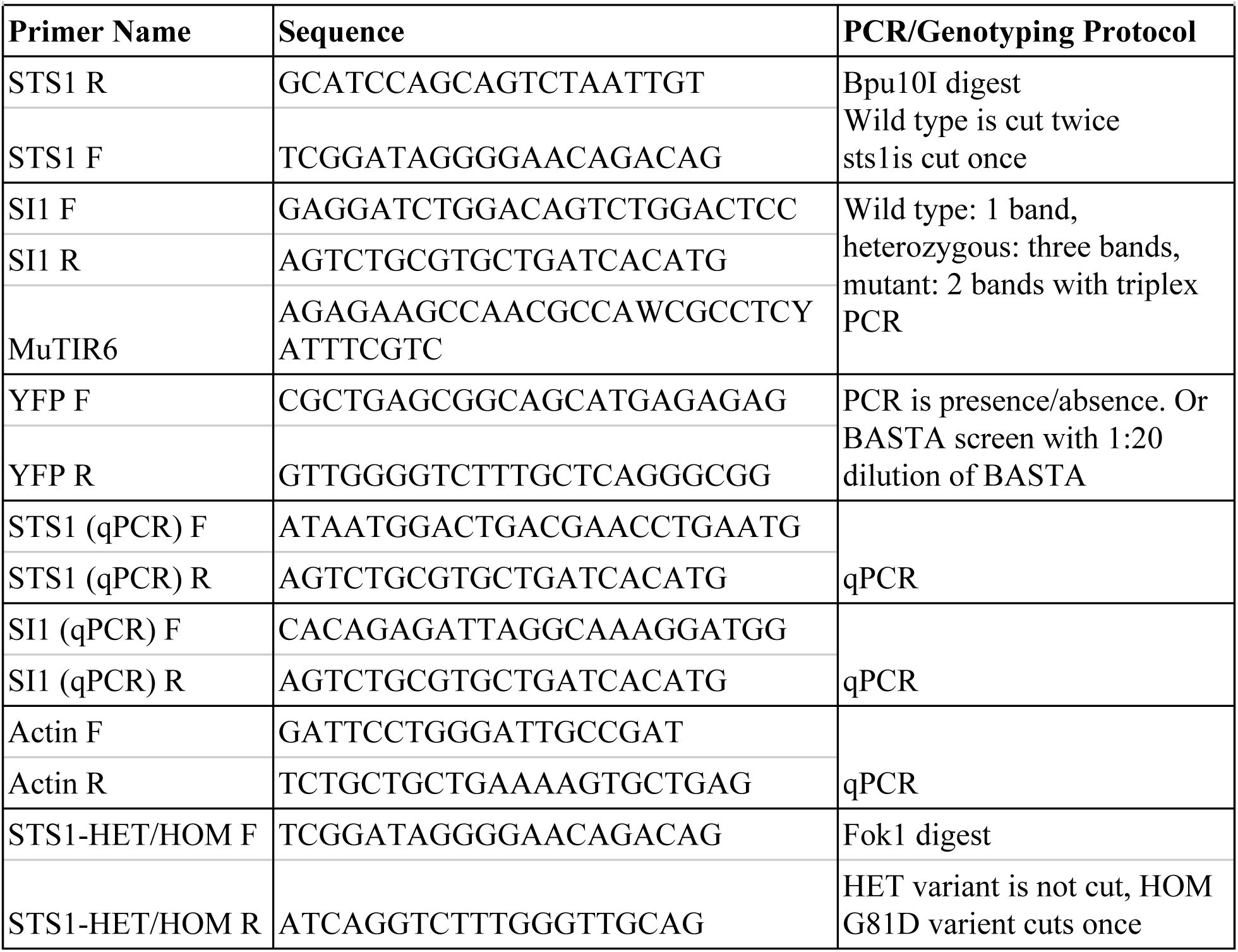
Primers for genotyping assays and RT-qPCR.

